# Farnesoid X receptor-dependent microbiome-bile acid signaling mediates obstructive sleep apnea-induced atherosclerosis

**DOI:** 10.64898/2026.03.31.715631

**Authors:** Jin Xue, Celeste Allaband, Simone Zuffa, Dan Zhou, Orit Poulsen, Jason Meadows, Daniel McDonald, Madison Ambre, Gail Ackermann, Amanda Birmingham, Jennifer Cao, Ipsita Mohanty, Pieter C. Dorrestein, Rob Knight, Gabriel G. Haddad

**Affiliations:** Department of Pediatrics, University of California San Diego, La Jolla, CA, USA; Skaggs School of Pharmacy and Pharmaceutical Sciences, University of California San Diego, La Jolla, CA, USA; Collaborative Mass Spectrometry Innovation Center, University of California San Diego, San Diego, CA, USA; Center for Microbiome Innovation, University of California, San Diego, La Jolla, CA, USA; Department of Computer Science and Engineering, University of California, San Diego, La Jolla, CA, USA; Shu Chien-Gene Lay Department of Bioengineering, University of California San Diego, La Jolla, CA, USA; Halıcıoğlu Data Science Institute, University of California San Diego, La Jolla, CA, USA; Hong Kong University of Science and Technology Jockey Club Institute for Advanced; Study, Hong Kong SAR, China; Department of Neuroscience, University of California San Diego, La Jolla, CA, USA; Rady Children’s Hospital, San Diego, CA, USA

**Keywords:** Atherosclerosis, FXR, gut microbiome-bile acid axis, microbial bile acid modifying enzymes

## Abstract

Intermittent hypoxia and hypercapnia (IHC), a hallmark of obstructive sleep apnea (OSA), accelerates atherosclerosis, yet the underlying mechanisms remain unclear. The gut microbiota and metabolites, specifically bile acids, change with IHC and thus the bile acid receptor farnesoid X receptor (FXR) might mediate IHC-induced atherosclerosis. In this study, *ApoE^-/-^* and *ApoE^-/-^ FXR^-/-^* mice were exposed to IHC or room air and fed with a high-fat, high-cholesterol diet for 10 weeks. Markers of atherosclerosis, fecal microbiome, and metabolome were then examined via Sudan IV staining, absolute abundance shotgun metagenomics, and untargeted liquid chromatography tandem mass spectrometry (LC-MS/MS). IHC markedly increased aortic atherosclerosis in *ApoE^-/-^*mice, an increase that was abolished by FXR deficiency. In addition, IHC reshaped gut microbial composition, promoting enrichment of bile acid–modifying taxa and increasing levels of microbial hydroxysteroid dehydrogenase (*hsdh*). The bile acid pool was also remodeled and associated with aortic atherosclerosis via FXR-dependent metabolic signals in *ApoE^-/-^* mice. Knockout of FXR disrupted microbiome shift under IHC and uncoupled microbial bile acid metabolism from vascular lesion development, thereby protecting against aortic atherosclerosis. These findings show that FXR has a central role in linking IHC, microbial bile acid metabolism, and cardiovascular pathology.

## INTRODUCTION

Obstructive sleep apnea (OSA) is a common sleep disorder caused by recurrent collapse of the upper airway, leading to oxygen desaturation and sleep fragmentation. OSA affects almost 1 billion people globally.^1^ In the United States, about 54 million people meet the criteria for OSA, 25-30% in men, 9-17% in women, and 2-3% in children.^2–4^ Importantly, the prevalence of OSA increases with age and obesity^3^ and clinical studies have shown that OSA increases morbidity and mortality from cardiovascular diseases, including atherosclerosis^5–8^. Given the widespread impact of OSA, there is a need for increased awareness, diagnosis, treatment, and prevention. However, the mechanisms underlying OSA-induced atherosclerosis remain elusive.

We have previously showed that intermittent hypoxia and hypercapnia (IHC), a hallmark of OSA pathophysiology, exacerbates atherosclerosis beyond the effect of high fat high cholesterol (HFHC) diet, which is associated with significant alterations in both the fecal microbiota and metabolome.^9–12^ Notably, germ-free (GF) *ApoE^-/-^*mice exhibited reduced atherosclerotic lesions under IHC/HFHC conditions compared to conventionally-reared specific pathogen free (SPF) *ApoE^-/-^*mice, identifying the gut microbiota as a possible causal factor in IHC/HFHC-induced atherogenesis.^13^

Interestingly, we found that bile acids were among the most dysregulated metabolites by IHC.^13^ Bile acids are synthesized from cholesterol, primarily in the liver but distant organs also contribute to the bile acid pool.^14^ It has been shown that Western diet is more atherogenic.^14^ Our previous atherosclerosis study on different diets revealed that high cholesterol, rather than high fat, is essential for IHC-induced atherosclerosis^13^, implying the importance of bile acid production from cholesterol. The gut microbiota encodes enzymes that can transform primary bile acids into secondary bile acids via deconjugation, dehydroxylation, oxidation/epimerization, desulfation, esterification, and amidation, which in turn influence host health by modulating metabolism, inflammation, and immunity.^15–17^ However, the contributions of microbiota-modulated bile acids in IHC-induced atherosclerosis have not been thoroughly investigated. In the host, bile acids act as signaling molecules and bind to receptors, such as nuclear farnesoid X receptor (FXR) and membrane bound Takeda G protein-coupled receptor (TGR5), which regulate various metabolic processes, including glucose and lipid metabolism, energy balance, inflammation as well as cardiovascular function.^15,16,18,19^ The role of FXR in atherosclerosis is multifaceted. Activation of FXR offers beneficial effects by improving plasma lipid profiles and reducing inflammation^20^ but can also negatively impact HDL (high-density lipoprotein) cholesterol levels which are inversely correlated with atherosclerosis risk^21^. Moreover, FXR regulates vascular tension and the unloading of cholesterol from foam cells, thereby affecting the progression of atherosclerosis^20^. Results from FXR deficiency studies have demonstrated protective effects. Knockout of the FXR gene has been shown to attenuate atherosclerosis in both *Ldlr^-/-^*^22^ and *ApoE^-/-^*mice^23^ under different experimental settings. Additionally, suppression of intestinal FXR signaling either by gene knockout or by pharmacological inhibition also showed decreased atherosclerosis.^24^ Hence, further research is warranted to fully understand the complex interplay between FXR, bile acids, and atherosclerosis and to determine the optimal strategies for targeting FXR in the treatment of cardiovascular disease.

In the present study, we investigate (1) the role of FXR in IHC-induced atherosclerosis; (2) the differentially modulated gut bacteria and metabolites under IHC and Air conditions in *ApoE^-/-^ FXR^-/-^* mice compared to *ApoE^-/-^* mice; (3) the relationship between FXR, gut microbiota, microbial-modified bile acids, and IHC-induced atherosclerosis. Our results enhance our understanding of the mechanisms mediating OSA-induced atherosclerosis and provide novel targets for better treatment and prevention of OSA-related cardiovascular complications.

## MATERIALS AND METHODS

### Animals and Diets

*ApoE*^-/-^ mice (Strain 002052) and *FXR*^-/-^ mice (Strain 007214), both on C57BL/6J background, were purchased from Jackson Laboratory (Bar Harbor, ME). Mice carrying deletion of both *FXR/Nr1h4* and *ApoE* genes (i.e., double KO of *FXR* and *ApoE*) were generated by in- house breeding of *ApoE*^-/-^ mice and *FXR*^-/-^ mice. *ApoE* and *FXR* deficiencies were confirmed by polymerase chain reactions (PCR). Nine to thirteen week old male mice were placed on a high fat and high cholesterol diet (HFHC) containing 1.3% cholesterol by weight and 42% fat by Kcal (TD.96121; Envigo-Teklad, Madison, WI), either under IHC or Air, for 10 weeks as previously described.^13^ The body weight of each mouse was measured and the food consumption of each mouse cage was recorded once per week. Fecal samples for microbiome and metabolome analysis were collected every two weeks and frozen at -80 °C until further processing. The collection time window between 9AM and 11AM (ZT3-ZT5) was chosen based on findings from a concomitant circadian study by our group, which identified ZT3–ZT5 as the period with the most pronounced microbiome composition differences between IHC and Air.^25^ All animal protocols were approved by the Animal Care Committee of the University of California San Diego and followed the Guide for the Care and Use of Laboratory Animals of the National Institutes of Health.

### Intermittent Hypoxia and Hypercapnia Treatment

Intermittent hypoxia and hypercapnia (IHC) were administered in a computer-controlled atmosphere chamber (OxyCycler, Reming Bioinstruments, Redfield, NY) as previously described.^9^ Each IHC cycle lasted 10 minutes with ∼4 minutes of 8% O_2_ and 8% CO_2_, followed by 1–2 minutes ramp intervals and ∼4 minutes of normoxia (21% O_2_) and normocapnia (0.5% CO_2_). The treatment period was 10 hours per day during the light cycle, for 10 weeks, starting at the same time when the diet was switched from regular chow to HFHC. Control mice were under room air (21% O_2_ and 0.5% CO_2_) and fed with the same HFHC.

### Quantification of Atherosclerotic Lesions

After 10 weeks of IHC and HFHC treatment, the mice were perfused with 4% paraformaldehyde. The entire aorta and the pulmonary arteries (PAs) were dissected and stained with Sudan IV. Atherosclerotic lesion areas were quantified by computer-assisted image analysis (ImageJ, NIH Image)^26^ as previously described^9^ and expressed as the percentage of Sudan IV positive area to the total vessel area. The aortic arch was consistently defined by cropping at a fixed distance from the bifurcation using Adobe Photoshop (Adobe Photoshop CS6, Adobe Systems Inc., San Jose, CA). All analyses were conducted in a blinded manner. Data are presented as mean ± SEM. Statistical significance was assessed by one-way ANOVA with Tukey’s post hoc test. An adjusted p < 0.05 was considered significant.

### Microbiome Analysis

#### Sample processing

Fecal pellet samples (∼0.03 - 0.04g) were collected in 1 mL Matrix Tubes (ThermoFisher, Waltham, MA, USA) for gDNA extraction.^27^ Extraction was performed using reagents from the MagMAX Microbiome Ultra Nucleic Acid Isolation Kit (ThermoFisher Scientific, Waltham, MA, USA) following a previously described high-throughput protocol^28^, minimizing reaction volumes, eliminating the need for the sample-to-bead plate transfer step, and reducing potential cross-contamination from vortexing during bead beating. The gDNA was quantified using the Quant-iT PicoGreen dsDNA Assay Kit (Invitrogen, Waltham, MA, USA) and normalized to 5 ng in 3.5 µL sterile water for library preparation, which was performed using a miniaturized adaptation of the KAPA HyperPlus Library Kit (Roche, Basel, Switzerland).^29^ The library preparation was quantified via a PicoGreen Assay (Invitrogen), and all samples were equal volume pooled, PCR cleaned (Qiagen), and size selected from 300 to 700 bp using a Pippin HT (Sage Sciences). Quality control was run on an Agilent 4200 Tapestation (Agilent, Santa Clara, CA, USA) to confirm expected library sizes after PCR cleanup and size selection. The equal volume pool was sequenced on an iSeq 100 (Illumina, San Diego, CA, USA). Utilizing the sample concentration and read counts per sample obtained from the iSeq 100 run, a normalized pooling value was calculated for each sample to optimize pooling efficiency to obtain more even read counts per sample during NovaSeq X Plus sequencing.^30^ After re-pooling the library with the iSeq-normalized pool values, samples were PCR cleaned and size selected (300–700 bp), and another round of quality control was performed using an Agilent tapestation. A synDNA spike in was used for absolute quantification of microbial abundances.^31^

#### Bioinformatic analysis

The resulting pool of PCR-cleaned, size-selected gDNA was sequenced on a NovaSeq X Plus (Illumina, San Diego, CA, USA) at the Institute for Genomic Medicine at the University of California, San Diego with a 25B flow cell and 2 × 150 bp chemistry. Raw BCL sequence read files were demultiplexed to per sample FASTQs and quality filtered.^32^ Adapter trimming was performed by fastp^33^ and the resulting FASTQ files were uploaded into Qiita^34^ ([study ID #15424](https://qiita.ucsd.edu/study/description/15424)). Within Qiita, mouse reads were filtered using GRCm39 and then processed using the Woltka^35^ pipeline (version 0.1.7) in paired-end mode with SHOGUN^36^ parameters. In brief, direct genome alignments were made against the “Web of Life” database^37^ (release 2), which contains 15,953 microbial genomes of bacteria and archaea. Paired-end sequence alignment was performed using the bowtie2^38^ (v2.5.4) aligner (using --very-sensitive -k 16 --np 1 --mp “1,1” --rdg “0,1” --rfg “0,1” --score-min “L,0,-0.05” --no-head --no-unal --no-exact-upfront --no-1mm-upfront) to map sequencing data to microbial reference genomes.^36^ Microbial genome IDs were considered operational genomic units (OGUs).^35^ The resultant count matrix was saved as a BIOM-format table.^39^

We applied a novel coverage dispersion filter called micov^40^ (https://github.com/biocore/micov) to identify the percent reference genome coverage. The counts matrix, OGU coverage information and lengths, and sample and synDNA input masses were provided to the pysyndna software (https://github.com/biocore/pysyndna). This tool implements and extends the calculations described in Zaramela, *et al.*^31^, producing for each sample the estimated cell counts per gram (i.e. absolute abundance) of sample for each OGU. OGUs with micov reference genome coverage of less than 1% - were excluded from calculations to reduce risk of false positives from short read multimapping. The absolute abundance BIOM file was converted to a QIIME2^41^ (v2025.4) artifact. Alpha diversity metrics (Faith’s PD^42^, Shannon index^43^) and the beta diversity phylogenetic robust center-log-ratio (rclr) metric (phylo-RPCA^44^) were calculated using unrarefied absolute abundance values. Greengenes2^45^ (v.2024.09) was used for taxonomy and phylogeny. Additionally, TEMPoral TEnsor Decomposition (TEMPTED)^46^, a beta diversity dimensionality reduction method that accounts for repeated sampling and uses time as a continuous variable, was used. Alpha diversity group differences were determined by non-parametric statistical testing, either Kruskal-Wallis with post-hoc Dunn’s test or two-sided Mann-Whitney-Wilcoxon with Holm-Bonferroni correction for multiple comparisons. Beta diversity differences were determined by pairwise PERMANOVA with post hoc Dunn’s test. Feature tables at the OGU level were used to examine differential abundance. Linear mixed-effects models (LME) were used via the statsmodels^47^ (0.14.2) package in Python to assess whether dependent variables of interest vary with fixed effects. Generally, OGU absolute abundance(cells/g) was used as a fixed effect and individual host subject identification was used to account for repeated measures. For example, the LME formulas were similar to “OGU(cells/g) ∼ timepoint + genotype_exposure + (1 | host_subject_id)”. If included, a star indicates an interaction term. The Benjamini-Hochberg procedure was applied to control for false discovery rate (FDR). Top and bottom differentially ranked features from phylo-RPCA and TEMPTED Axis 2 were also used to identify OGU of interest.

#### Gene searches

Gene searches were specifically performed for bile acid modification enzymes (*hsdh*, *bai*, *bsh*, *assT*). For each gene of interest, faa files of all matches were acquired from Uniprot^48^ and pooled; amino acid sequences with at least 60% similar identity were then clustered via USEARCH^49^into a unique database. Then, metagenomic reads from each sample were aligned and matched to the genes of interest against the respective USEARCH clustered databases using Diamond blastx^50^. Output data frames were further filtered for matches with > 40 bp of alignment length, > 50% query alignment ratio, and e-value < 0.00001 before plotting.

### Metabolomics analysis

#### Sample processing

Samples were acquired via untargeted metabolomics using a Vanquish UHPLC system coupled to a Q-Exactive Orbitrap mass spectrometer (Thermo Fisher Scientific), as previously described.^51^ The chromatographic separation was performed on a polar C18 column (Kinetex C18, 100 x 2.1 mm, 2.6 μm particle size, 100A pore size – Phenomenex, Torrance, USA), and the mobile phase consisted of H_2_O (solvent A), and acetonitrile (solvent B), both acidified with 0.1% formic acid. The LC method consisted of 0-0.5 min 5% B, 0.5-1.1 min 5-25% B, 1.1-7.5 min 25-40% B, 7.5-8.5 min 40-100% followed by a 1.5 min washout phase at 100% B, and a 2.0 min re-equilibration phase at 5% B. The flow rate was set at 0.5 mL/min, the injection volume was fixed at 5 μL, and the column temperature was set at 40 °C. Data-dependent acquisition (DDA) of MS/MS spectra was performed in the positive ionization mode. Electrospray ionization (ESI) parameters were set as: 52.5 AU sheath gas flow, 13.75 AU auxiliary gas flow, 2.7 AU spare gas flow, and 400 °C auxiliary gas temperature; the spray voltage was set to 3.5 kV and the inlet capillary to 320 °C and 50 V S-lens level was applied. MS scan range was set to 150-1500 *m/z* with a resolution of 35,000 with one micro-scan. The maximum ion injection time was set to 100 ms with an automated gain control (AGC) target of 1.0E6. Up to 5 MS/MS spectra per MS1 survey scan were recorded in DDA mode with a resolution of 17,500 with one micro-scan. The maximum ion injection time for MS/MS scans was set to 150 ms with an AGC target of 5E5 ions. The MS/MS precursor isolation window was set to 1 *m/z*. The normalized collision energy was set to a stepwise increase from 25 to 40 to 60 with z = 1 as the default charge state. MS/MS scans were triggered at the apex of chromatographic peaks within 2 to 5 s from their first occurrence.

#### Data processing

Generated .raw files were converted into open-access format .mzML using ProteoWizard MSConvert^52^ and deposited under the accession code [MSV000098881](https://massive.ucsd.edu/ProteoSAFe/dataset.jsp?task=915b919d472b4b5fa2bf6e8c62b6770a) in GNPS/MassIVE. Feature extraction was performed using MZmine^53^ (version 4.2) via batch processing. In short, MS1 and MS2 noise levels were set to 5E4 and 1E3. Five minimum scans and a minimum absolute height of 1E5 were used for the chromatogram builder. Smoothing, local minimum resolver, 13C isotope filter, and isotopic peak finder were applied before the join aligner, which had a 0.20 retention time tolerance. Features present in fewer than 2 samples were removed before gap filling (feature finder). Ion identity networking and metaCorrelate were applied before extrapolating the final .mgf and feature table files. Feature-based molecular networking^54^ was performed in GNPS2^55^ using a minimum of five matching fragment ions, a tolerance of 0.02, and cosine score similarity > 0.7. Generated annotations are level 2 according to the Metabolomics Standard Initiative.^56^ CANOPUS^57^ was used to predict molecular classes of the MS/MS spectra via SIRIUS^58^ (version 5.8).

#### Data analysis

Data were imported in R 4.5.2 (R Foundation for Statistical Computing, Vienna, Austria) for downstream data analysis. A cleaning step (blank substraction) was performed to remove features whose intensity in samples was not at least 5 times that observed in blanks. Principal component analysis (PCA) and partial least-squares discriminant analysis (PLS-DA) were performed using mixOmics^59^ (version 6.34.0) after RCLR transformation using vegan^60^ (version 2.7-2). After initial data overview, the first timepoint was removed from downstream analysis as it was collected at baseline when the animals were on a standard chow diet. Stratification based on genotype or gas exposure was used to extract influenced features. Variable importance (VIP) scores were considered relevant for the discrimination when > 1. PLS-DA model performances were evaluated using a 10-fold cross-validation and 999 permutations. A volcano plot was used to visualize linear regression output of features associated with percentage of atherosclerosis in the aorta. The lmerTest^61^ (version 3.2-0) package was used for linear mixed effect models. Upset plots were generated using UpSetR^62^ (version 1.4). Presence of the bile acid diagnostic ions^63^ was validated via MassQL^64^ queries using the dedicated MetaboApp^65^ (https://massqlpostmn.gnps2.org/).

### Multi-omics analysis

Joint-RPCA is an unsupervised method used to agnostically determine co-occurrences between microbiome and metabolome data.^66,67^ Samples from baseline (TP1) were excluded due to a different diet. In short, the unrarefied absolute count tables for each method after previously described filtration methods were RCLR transformed. RCLR of the raw observed values were only computed on the non-zero entries and then averaged and optimized. This helps to account for the sparsity of the microbiome data. Absolute count data is not compositional. The shared samples of all input matrices were used to estimate a shared matrix. The estimated shared matrix and the matrix of shared eigenvalues across all input matrices were recalculated at each iteration to ensure consistency. Minimization was performed across iterations by gradient descent. Cross-validation of the reconstruction was performed in order to prevent overfitting of the joint factorization. Co-occurrences of features across the two input matrices were calculated from the final estimated matrices.

### Data availability

Untargeted LC-MS/MS data generated for this study are publicly available at GNPS/MassIVE under the accession code [MSV000098881](https://massive.ucsd.edu/ProteoSAFe/dataset.jsp?task=915b919d472b4b5fa2bf6e8c62b6770a). The associated FBMN job is publicly available at GNPS2: https://gnps2.org/status?task=66819d0119814501806c2266f2d67e10. Metagenomics sequencing data is available at EBI/ENA via the following accession codes: PRJEB96078, ERP178828. Additional information and processing pipelines are available in Qiita under [Study ID #15424](https://qiita.ucsd.edu/).

### Code Availability

Code used to analyze and generate the figures for the metagenomics analyses is located at https://github.com/knightlab-analyses/haddad_apoe_fxr_KO_abs_quant, while code relative to the metabolomics analyses is located at https://github.com/simonezuffa/Manuscript_FXR_IHC.

## RESULTS

### FXR deficiency abolished IHC-induced atherosclerosis in the aorta

To investigate if FXR mediates IHC-induced atherosclerosis, we exposed *ApoE^-/-^FXR^-/-^* mice and *ApoE^-/-^* mice to either IHC or Air, both fed with a HFHC diet for 10 weeks. Lesion data revealed that IHC promoted the development of atherosclerosis in the aorta and pulmonary artery (PA) of *ApoE^-/-^* mice, consistent with our prior findings^13^ (*ApoE^-/-^*mice, Aorta: IHC 16.7 ± 1.46% vs Air 10.3 ± 0.26%, p < 0.05; PA: IHC 25.3 ± 2.12% vs Air 8.8 ± 1.12%, p < 0.001) (**Figure 1**). Intriguingly, knockout of FXR diminished lesions in the aorta and aortic arch under IHC, which were similar to lesion levels under Air, suggesting that FXR receptor is a key mediator in IHC-induced atherosclerosis (Aorta: IHC, *ApoE^-/-^FXR^-/-^* 7.0 ± 1.70% vs *ApoE^-/-^* 16.7 ± 1.46%, p < 0.001; vs Air *ApoE^-/-^ FXR^-/-^* 6.9 ± 2.06%, p > 0.05. Aortic arch: IHC, *ApoE^-/-^ FXR^-/-^* 15.9 ± 3.05% vs *ApoE^-/-^*26.4 ± 1.68%, p < 0.05; vs Air *ApoE^-/-^ FXR^-/-^* 13.2 ± 2.49%, p > 0.05) (**Figure 1**). In contrast to the aorta, IHC-induced atherosclerosis was still observed in *ApoE^-/-^ FXR^-/-^* mice in the PA, suggesting that development of atherosclerosis might be influenced by different mechanisms in the PA (*ApoE^-/-^ FXR^-/-^* mice, PA: IHC 19.4 ± 4.00% vs Air 9.2 ± 2.49%, p < 0.05) (**Figure 1**).

**Figure 1.**
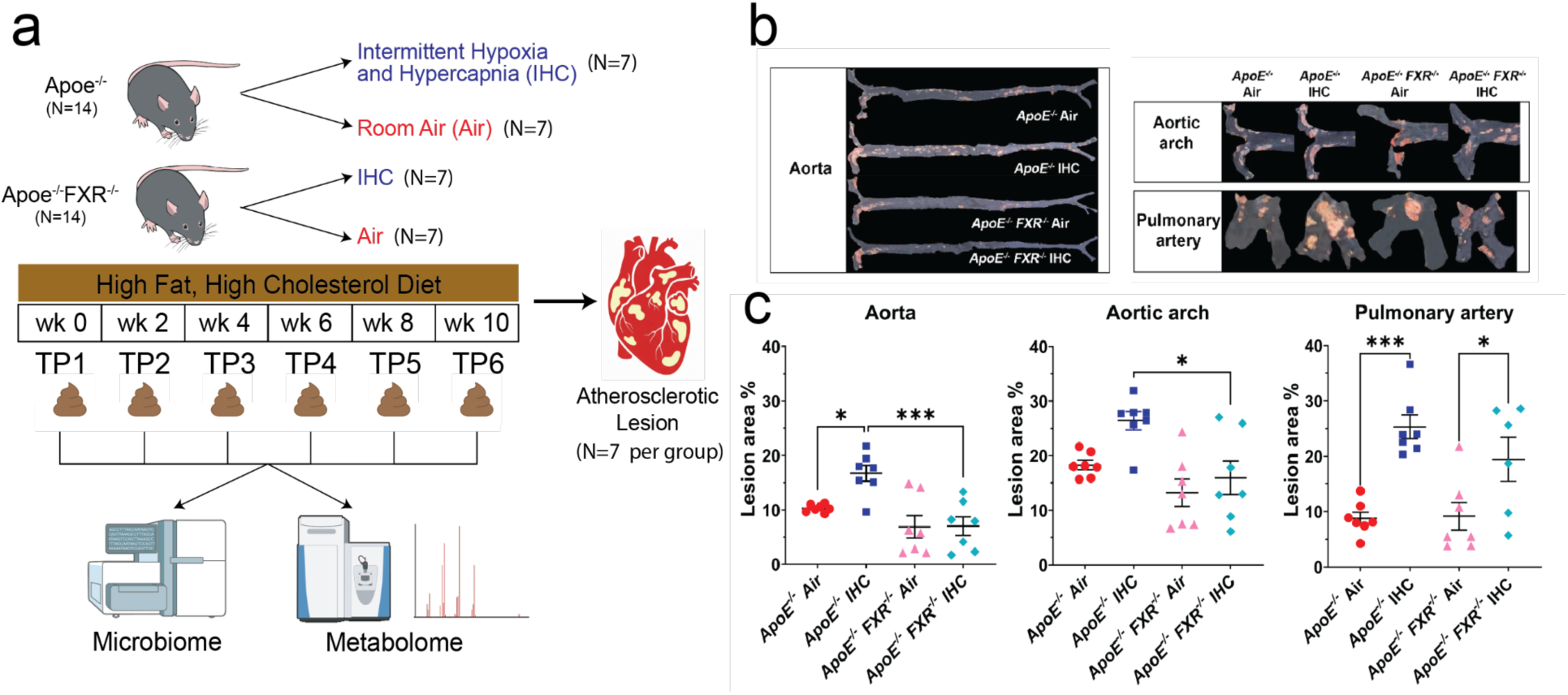
FXR Knockout Attenuates Atherosclerotic Formation in the Aorta. **a)** Experimental design. *ApoE^-/-^*and *ApoE^-/-^ FXR^-/-^* male mice at the age of 9-13 weeks old were exposed to either IHC or Air, both fed with HFHC diet for 10 weeks (N=7 for each group). Aorta and PA were harvested at the end of 10-week treatment for atherosclerotic lesion analysis. Fecal samples were collected every 2 weeks for time course analyses of microbiome and metabolome. Body weight and food intake were recorded every week. Images created using NIH BioArt (https://bioart.niaid.nih.gov/) and Google Gemini/NanoBanana2. **b)** Representative images of atherosclerosis stained with Sudan IV. And **c)** the en-face lesions were quantified as the percentage of lesion area in the total area of the blood vessel examined. IHC induced atherosclerosis was significantly reduced in the aorta and aortic arch of *ApoE^-/-^ FXR^-/-^* mice as compared to *ApoE^-/-^* mice. No significant difference was detected between *ApoE^-/-^* and *ApoE^-/-^ FXR^-/-^* mice under Air. Data are presented as means ± SEM. One-way ANOVA followed by Tukey’s multiple comparison test. Significance: * p < 0.05; *** p < 0.001.

### FXR deficiency abolished IHC-induced changes in the gut microbiome composition

We generated absolute abundance data from shotgun metagenomics and found that the total number of detected bacterial cells per gram of feces had a borderline significant negative correlation (linear regression: R^2^ = 0.14, p = 0.053) with aortic lesions at timepoint 6 (TP6) (**Figure S1a**). When examined by individual genotype and exposure group, Air groups tended to have a neutral to mildly positive correlation (slope) and IHC groups from both genotypes had a negative correlation between total cell count and aorta lesion size (**Figure 2a**). This indicated that under IHC conditions, reduced overall bacterial abundance was associated with increased aortic lesions. However, the total number of detected bacterial cells per gram of feces was not significantly different between exposure types for both genotypes over the course of the experiment (linear mixed effects model (LME) p>0.05) (**Figure S1b**). There were also no significant differences in absolute abundance between exposure types for either genotype for any of the alpha diversity metrics examined (LME p>0.05) (**Figure S1 c-e**). We generally found increasing total bacterial cell count appeared to be linked to increased phylogenetic diversity as expected, but correlations were weak: *ApoE^-/-^*Air (slope=+3.69×10^-8^), *ApoE^-/-^ FXR^-/-^* Air (-0.44×10^-8^), *ApoE^-/-^*IHC (slope= -2.93×10^-8^), *ApoE^-/-^ FXR^-/-^* IHC (slope=+6.21×10^-8^) (**Figure 2b**). Collectively, the lack of significant differences in absolute abundance and its weak correlation with alpha diversity metrics suggests that total microbial density was not a primary factor influencing the genotype- or exposure-specific outcomes in this model.

**Figure 2.**
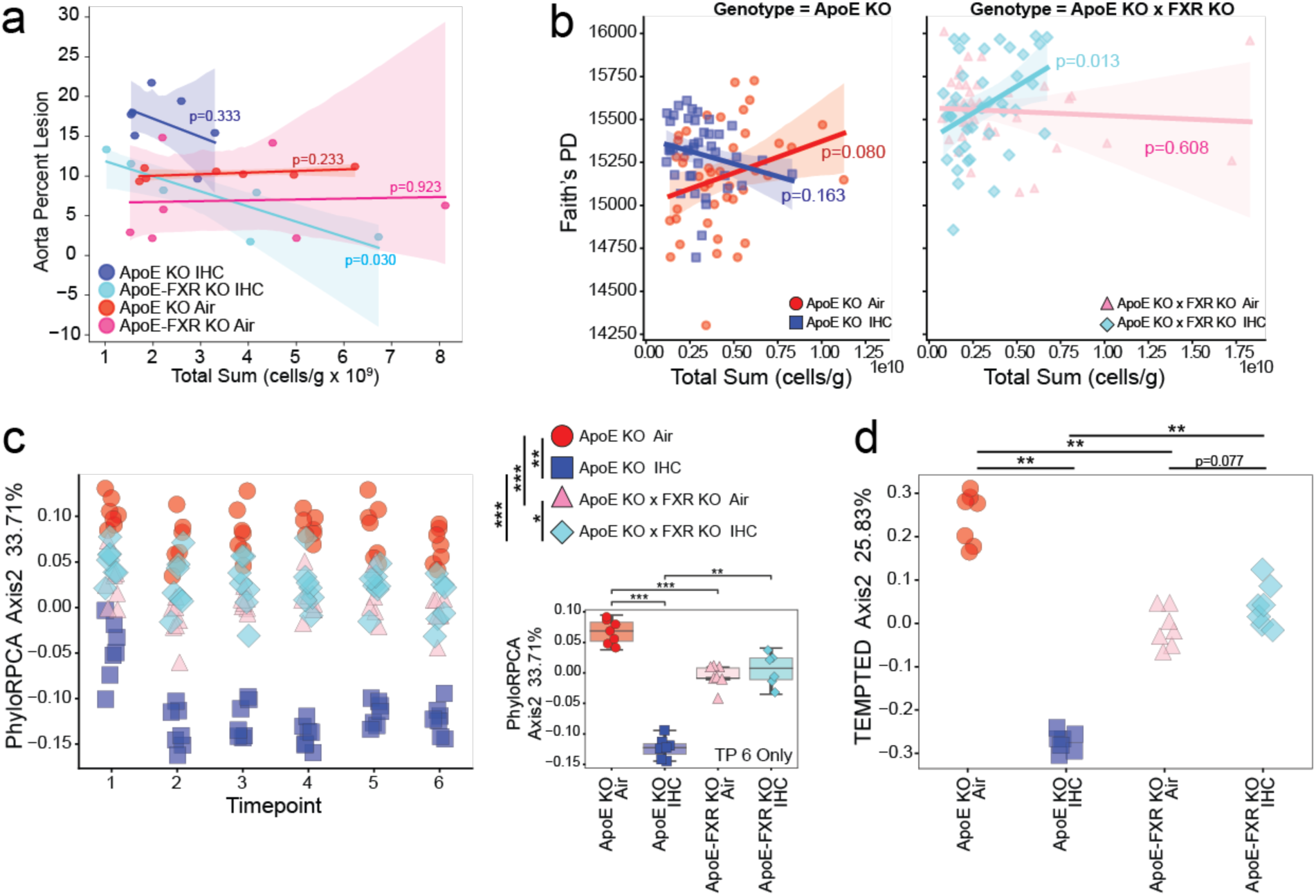
Absolute Abundance Metagenomic Identifies FXR Deficiency Protection against IHC-induced microbiome modulation. **a)** Absolute quantification (cells/g) comparison to percent lesions in aorta by genotype and exposure at the final timepoint only. **b)** Absolute abundance (cells/g) comparison to alpha diversity Faith’s PD metric by genotype and exposure across all timepoints. **c)** Beta diversity metric phylo-RPCA Axis 2 across all timepoints; significance determined by pairwise PERMANOVA with post-hoc Dunn’s test. Subset boxplot/stripplot is for TP6 only; significance determined by two-sided Mann-Whitney-Wilcoxon with Holm-Bonferroni correction. And **d)** longitudinal beta diversity metric TEMPTED Axis 2 by genotype and exposure. Significance determined by pairwise PERMANOVA with post-hoc Dunn’s test. Shaded regions for a and b indicate 95% confidence interval. Notation: * p < 0.05, ** p < 0.01, ***p < 0.001.

Microbiome composition, as measured by phylogenetic rclr-based beta diversity metric phylo-RPCA, revealed the strongest impact on the microbiome was genotype, which separated groups along Axis 1 (**Figure S1f**). The second most important effect on composition was exposure type, which separated groups along Axis 2 (**Figure 2c**). IHC and control (room air) exposures were significantly different in *ApoE^-/-^*mice (PERMANOVA, pseudoF = 586, p_adj_ = 0.001), an effect that was reduced in *ApoE^-/-^ FXR^-/-^* mice (PERMANOVA, pseudoF = 12, p_adj_ = 0.020). This was confirmed using longitudinal rclr beta diversity metric TEMPTED, where there were significant differences between IHC and room air controls for *ApoE^-/-^* mice (PERMANOVA, pseudoF = 351, p_adj_ = 0.003), which was not present for *ApoE^-/-^ FXR^-/-^* mice (PERMANOVA, pseudoF = 4, p_adj_ = 0.077) (**Figure 2d, Figure S1g**). In summary, the knockout of FXR ameliorated the shift that IHC exposure induced on the gut microbiome.

### OGU associated with *ApoE^-/-^* IHC mice with highest aortic lesions

Several different metrics for differential abundance were used to create a short list of specific OGU of interest. Linear mixed effects (LME) models, as well as the top and bottom differentially ranked features from phylo-RPCA Axis 2, and TEMPTED Axis 2 were employed. After evaluating overlap between the three methods, we highlighted specific OGU with the clearest patterns. We were especially interested in the changes that occurred in the latter half of the study and appeared to respond to increasing disease severity, especially those that were significantly different at the last timepoint (TP6). *Ruthenibacterium lactatiformans* (**Figure 3a**), *Sutterella wadsworthensis* (**Figure 3b**), *Faecousia sp000434635* (**Figure 3c**)*, Laedolimicola sp.* (**Figure 3d**), and *Gemmiger variabilis* (formerly *Subdoligranulum variabile*) (**Figure 3e**) all had elevated levels at the latter stages of the study primarily in *ApoE^-/-^* IHC mice, the group with the most atherosclerotic lesions. *Ruthenibacterium lactatiformans* responded particularly robustly at the final timepoints for *ApoE^-/-^* IHC mice. In contrast, *Intestinibacillus massiliensis* (**Figure 3f**) had decreased levels at the latter stages of the experiment in *ApoE^-/-^*IHC mice compared to the other groups.

**Figure 3.**
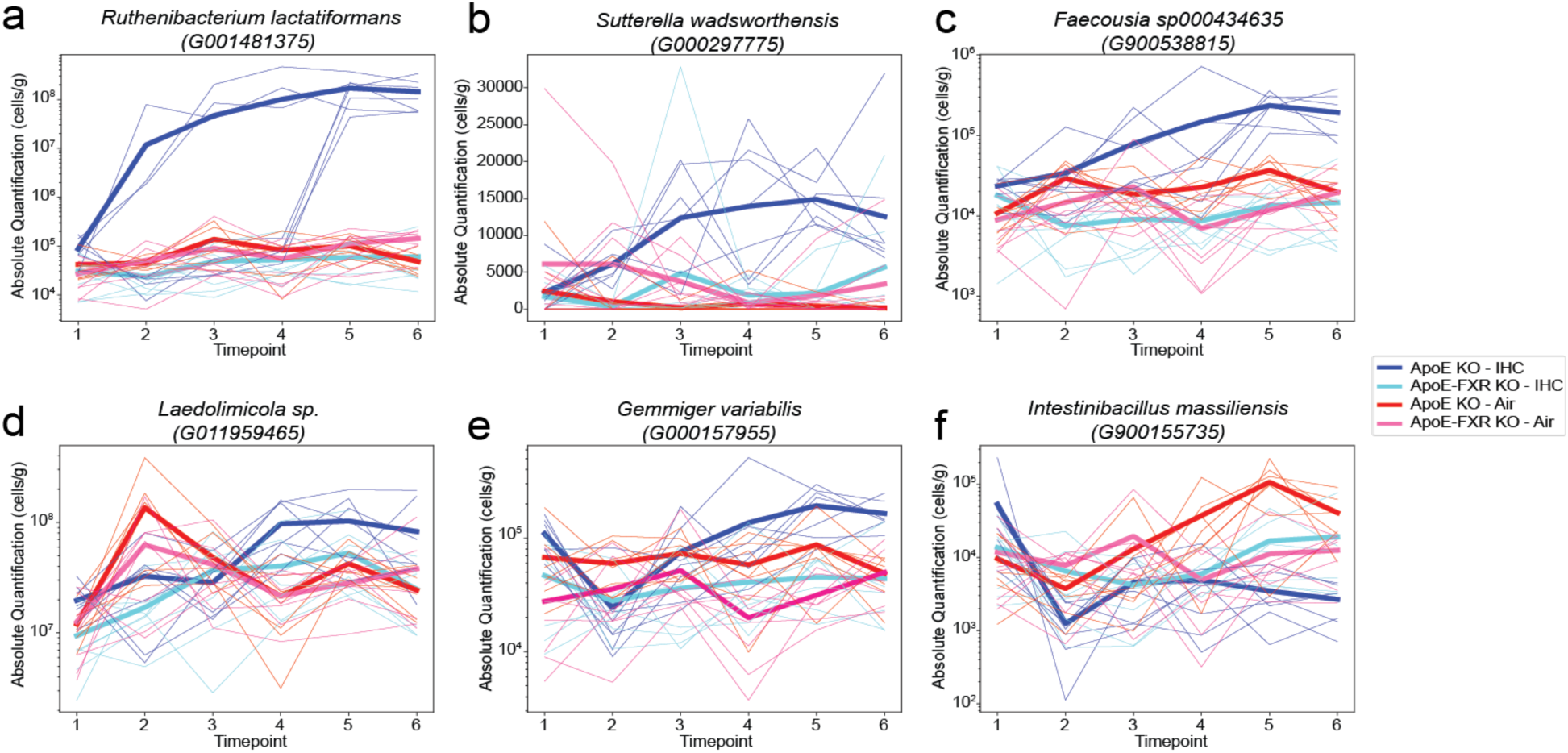
Microbial OGU Associated with ApoE^-/-^ IHC Mice with High Aortic Lesions, Especially at TP6, Using absolute abundance (cells/g). Highlighted OGU of interest that are differentially abundant in *ApoE^-/-^* IHC mice. All plots show groupings by both genotype and exposure. **a)** Lineplot of *Ruthenibacterium lactatiformans [Oscillospiraceae]*. *Oscillospiraceae* was formerly a subset of *Ruminococcaceae.* b) Lineplot of *Sutterella wadsworthensis [Sutterellaceae]*. **c)** Lineplot of *Faecousia sp000434635* (OGU G900538815) [*Oscillospiraceae*]. **d)** Lineplot of *Laedolimicola sp.* (OGU G011959465) [*Lachnospiraceae*]. **e)** Lineplot of *Gemmiger variabilis [Oscillospiraceae]*. And **f)** lineplot of *Intestinibacillus massiliensis [Erysipelotrichaceae]*. For all plots, thicker lines indicate group mean and thinner lines indicate individual mice.

In our previous work using 16S rRNA sequencing^12,13,25^, we observed members of *Akkermansiaceae* to respond to disease progression and to be significantly different between IHC and controls by the end of the study, which held true also for this study (**Figure S2a**). For *ApoE^-/-^* mice, *Akkermansiaceae* cell counts were lower in the mice under IHC conditions by the end of the study compared to controls. Similar to *Akkermansiaceae*, members of *Muribaculaceae* have been implicated in maintenance of the mucous layer and gut barrier integrity^68,69^ and were found to be lowest in *ApoE^-/-^* IHC mice with the greatest lesions in the current study, consistent with our previous finding. *Ruminococcaceae* and *Lachnospiraceae* are microbial families whose members are known to have bile acid modification genes.^70,71^ In this study, *Ruminococcaceae* showed patterns of interest, whereas *Lachnospiraceae* did not (**Figure S2c-d**). Additionally, the genus *Blautia A* (141780) is known for its ability to modify bile acids with species that have both *bsh* and *hsdh* genes^72,73^ and was one of the top differentially abundant genera with elevated amounts under *ApoE^-/-^* Air conditions for the last three timepoints (**Figure S2e**). Finally, the genus *Parasutterella* was also elevated in *ApoE^-/-^* IHC mice in the latter stages of the study and appeared to increase concomitantly with the lesion progression over time (**Figure S2f**).

### FXR KO protects from bile acid changes associated with atherosclerosis development

Untargeted metabolomics analysis of the fecal samples revealed an effect of the diet intervention (PERMANOVA, F = 19.79, p < 0.001), as samples collected at baseline were not on a HFHC diet, in conjunction with differences driven by both the genotype (PERMANOVA, F = 17.67, p < 0.001) and gas exposure (PERMANOVA, F = 4.83, p < 0.001) (**Figure 4a**). To investigate the effect of gas exposure (Air vs IHC), PLS-DA models stratified by genotype were created, excluding samples collected at baseline. Both models presented a strong classification performance (CER in *ApoE^-/-^* = 0.001, CER in *ApoE^-/-^ FXR^-/-^* = 0.09), with 2,140 and 2,137 metabolic features either enriched or depleted in response to IHC in the respective cohorts (**Table S1** and **Table S2**).

**Figure 4.**
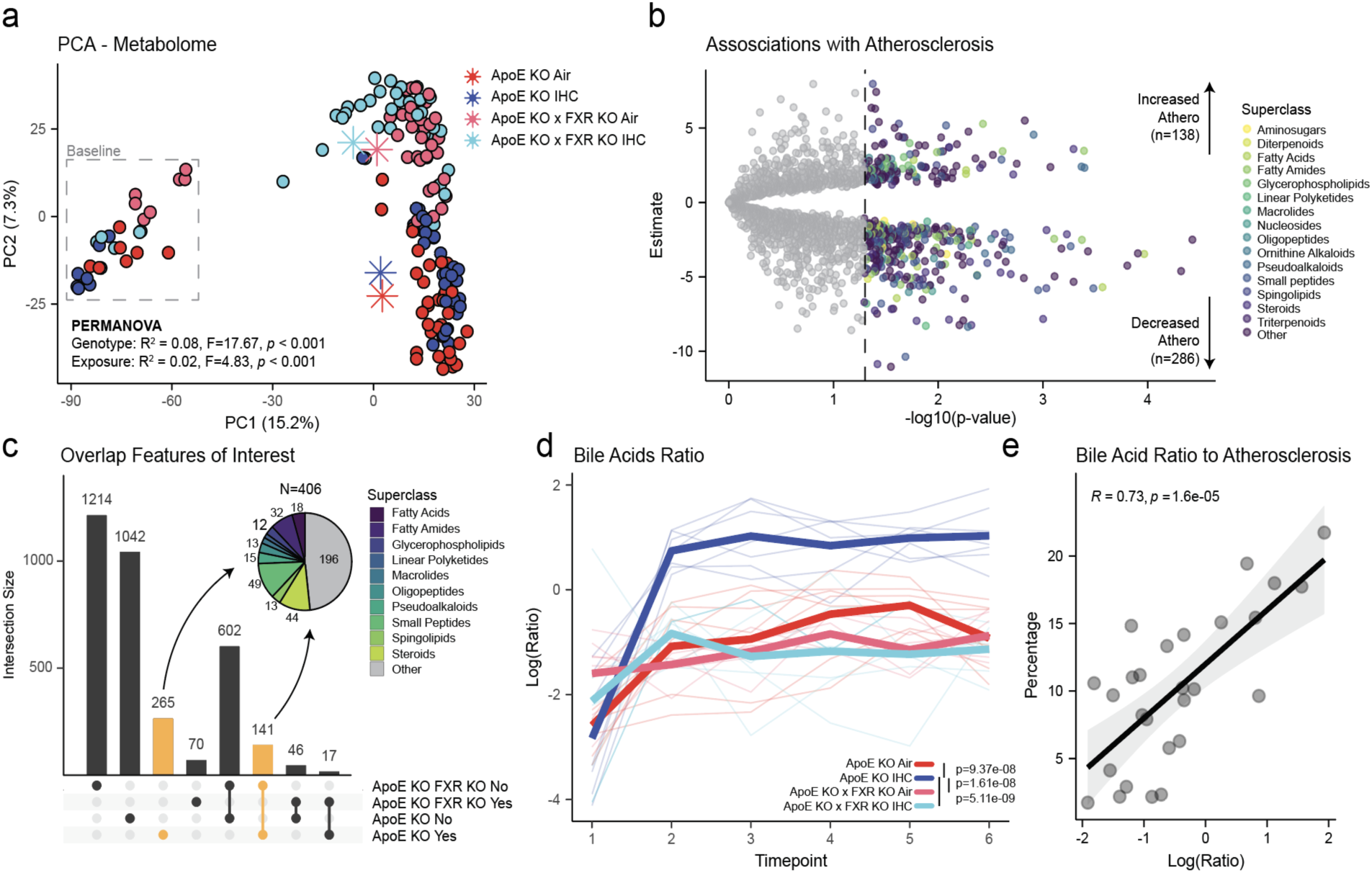
FXR KO Protects from Fecal Metabolic Changes Associated with Atherosclerosis. **a**) PCA of rclr transformed fecal metabolic profiles. PERMANOVA identified significant effects of diet intervention (F = 19.79, p < 0.001), genotype (F = 17.67, p < 0.001), and gas exposure (F = 4.83, p < 0.001). Asterisks represent group centroids. **b**) Volcano plot of features associated with atherosclerotic lesions (n=424) in the ApoE^-/-^ model. Estimates were calculated via linear regressions between the rclr transformed metabolite abundances at the last timepoint and the atherosclerotic lesions in the aorta. Only metabolic features that were significantly altered by gas exposure were investigated (VIP > 1 from PLS-DA model). **c**) Upset plot of significant features obtained from PLS-DA models based on gas exposure. Yes indicates correlation (positive or negative) to percentage of atherosclerosis lesions in the respective cohorts. No indicates no correlation. **d**) Line plot of natural log ratio of bile acids of interest influenced by IHC and correlating with atherosclerosis. The numerator consists of the sum of increased bile acids of interest (n=10) while the denominator for the decreased ones (n=42). Significance tested via linear mixed effect model with mouse id as random effect. And **e**) linear regression between bile acid ratio at the last timepoint (TP6) and aortic atherosclerosis in all animals (both genotypes included).

Given our interest in metabolites driving atherosclerosis, we then focused on fecal samples collected at the last timepoint from *ApoE^-/-^* mice only to test associations between rclr transformed abundances of the metabolites differentially modulated by gas exposure and atherosclerosis lesions in the aorta (**Table S3**). We then used a volcano plot to highlight 424 features either positively or negatively associated with atherosclerosis and their CANOPUS^57^ predicted superclasses (**Figure 4b**). After repeating the same analysis for the *ApoE^-/-^ FXR^-/-^* mice (**Table S4**), we extracted features either altered in the *ApoE^-/-^*mice only or altered in both genotypes but associated with atherosclerosis only in the *ApoE^-/-^*mice (**Figure 4c**). Interestingly, 265 features were found to be affected by gas exposure and correlated to atherosclerosis in the *ApoE^-/-^* mice only, while 141 features were affected in both genotypes but in the *ApoE^-/-^ FXR^-/-^* did not correlate with atherosclerosis. CANOPUS superclass predictions of the associated MS/MS spectra suggested that the majority of these features were small peptides, steroids, fatty amides, and fatty acids and conjugates. Given our previous studies^12,13,25^ and the investigation of FXR role, we focused on features annotated as bile acids. Out of the 406 features of interest, we identified 52 putative bile acids via spectral library annotation, CANOPUS prediction, or bile acid diagnostic ions. Next, we generated the log ratios for these bile acids and plotted it over the course of treatment (**Figure 4d**). The numerator comprised the sum of bile acids increased under IHC (n = 10), whereas the denominator comprised the sum of bile acids that were depleted (n=42). Linear mixed effect models confirmed the significance difference of *ApoE^-/-^* IHC group from the others (p < 1.85e-09). Interestingly, the differential ratio was established immediately after 2 weeks of intervention in the *ApoE^-/-^* IHC and remained stable throughout the duration of the study. The major change during this period was in diet (standard chow to HFHC) and gas exposure (standard air to IHC). Finally, the identified ratio showed a strong correlation (Pearson, R^2^ = 0.73, p = 1.6e-05) to aortic atherosclerosis at the last timepoint (TP6) (**Figure 4e**).

### Prevalence of bacterial *hsdh* positively correlated with atherosclerosis

Members of the gut microbiome encode genes involved in bile acid modification, such as *bsh* (bile salt hydrolase), *bai* (bile acid inducible), and *hsdh* (hydroxysteroid dehydrogenase). While in some cases *bai* genes are grouped with *hsdh* genes, we have kept them separately here. For example, *hsdh* genes encode enzymes that can add or remove hydroxyl groups, such as converting CDCA to UDCA or CA to UCA. These changes can impact FXR signaling, resulting in changes to host physiology, lipid homeostasis, and atherosclerotic disease progression.^74^ We observed a difference between genotype and exposure groups in detected prevalence of the *hsdh* gene over time (LME: IHC vs Air *ApoE*^-/-^ p <0.001, IHC vs Air *ApoE*^-/-^ *FXR*^-/-^ p = 0.128, Air *ApoE*^-/-^*FXR*^-/-^ vs *ApoE*^-/-^ p = 0.002, IHC *ApoE*^-/-^ *FXR*^-/-^ vs *ApoE*^-/-^ p < 0.001) (**Figure 5a**), including the final timepoint (2MWW-HB: *ApoE*^-/-^ IHC vs Air p < 0.001, *ApoE*^-/-^ *FXR*^-/-^ IHC vs Air p = 0.837, Air *ApoE*^-/-^ *FXR*^-/-^ vs *ApoE*^-/-^ p = 0.001, IHC *ApoE*^-/-^ *FXR*^-/-^ vs *ApoE*^-/-^ p = 0.022) (**Figure 5b**). Moreover, *hsdh* gene prevalence was positively correlated with aortic lesion prevalence in *ApoE*^-/-^ mice (Pearson, R^2^ = 0.59, p = 0.0013) (**Figure 5c**) but not in *ApoE*^-/-^ *FXR*^-/-^ mice (Pearson, R^2^ = 0.10, p = 0.3) (**Figure 5c**). In this study, *bai* genes were not different between groups (**Figure S3 a-b**), and *bsh* genes were only altered between IHC and control groups for *ApoE*^-/-^ *FXR*^-/-^ mice, which presented low levels of aortic atherosclerosis (**Figure S3 c-d**). Overall, this indicates that *hsdh* might play a pivotal role in both the balance of the bile acid pool, which affects FXR signaling, and atherosclerotic lesion formation.

**Figure 5.**
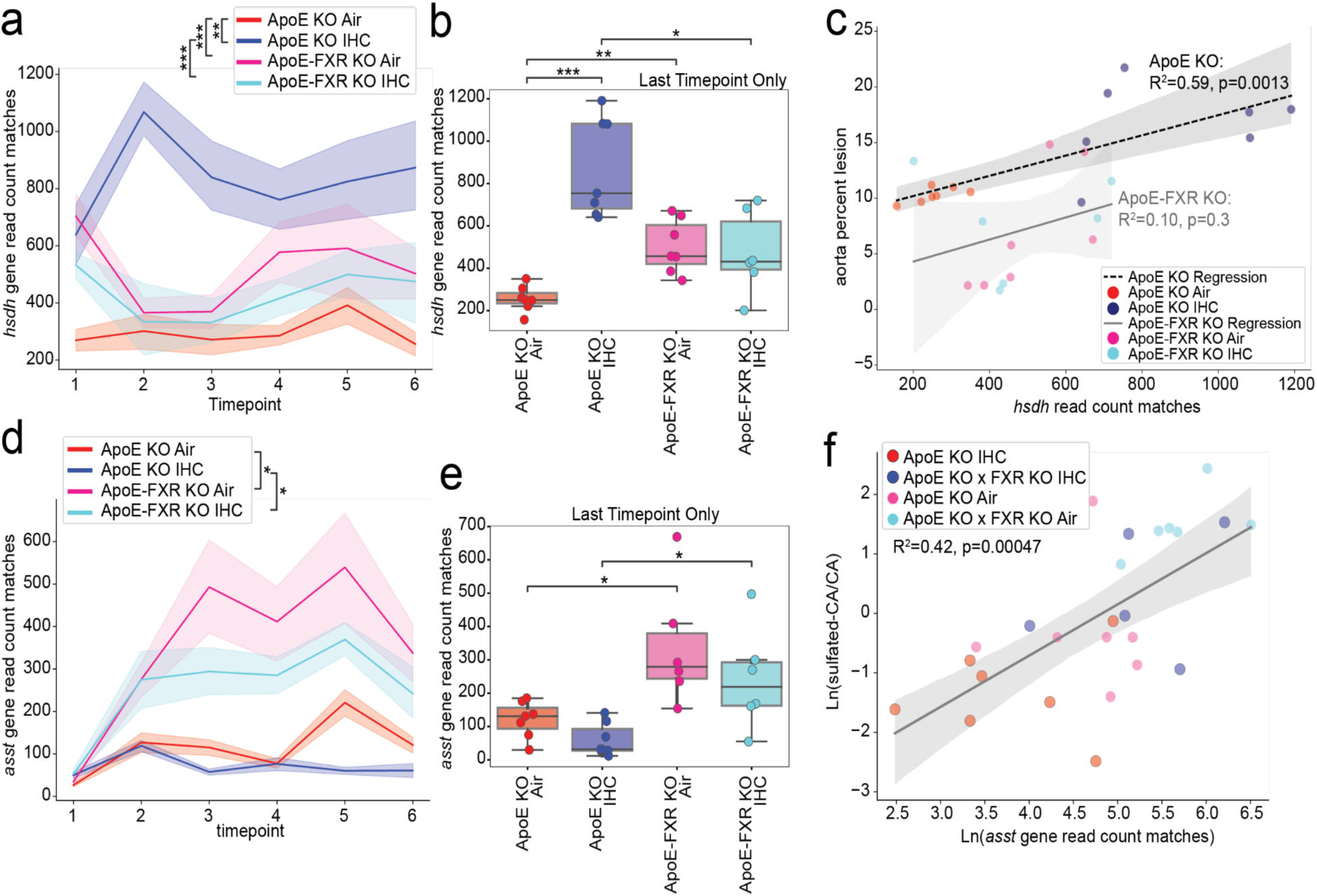
Bile Acid Modification Enzymes - hydroxysteroid dehydrogenase (hsdh) and aryl-sulfate sulfotransferase (*asst*) impact atherosclerosis and bile acid pool. **a)** Lineplot of *hsdh* read count matches by genotype and exposure over time; shaded region indicates standard error of the mean (SEM). **b)** Box-scatterplot of *hsdh* read count matches by genotype and exposure at the last timepoint only (TP6). **c)** Lmplot (linear regression) of *hsdh* read count matches from TP6 only compared to aorta lesions. **d)** Lineplot of *assT* read count matches by genotype and exposure over time; shaded region indicates SEM. **e)** Box-scatterplot of *asst* read count matches by genotype and exposure at TP6. And **f)** lmplot of the natural log of *asst* read count matches compared to the natural log of peak absorbance of putative sulfated cholic acid over putative cholic acid from TP6. Significance for a, d is determined by LME and accounts for repeated measures. Significance for b, e boxplots determined by two-sided Mann-Whitney-Wilcoxon with Holm-Bonferroni correction. Significance for lmplots determined by Pearson correlation. Notation: * p < 0.05, ** p < 0.01, *** p < 0.001.

Furthermore, bacterial *asst* (aryl-sulfate sulfotransferases) genes have been shown to transfer sulfate groups to phenols or other steroid compounds, like bile acids.^75^ However, the complete mechanism of *asst* enzymes, including substrates, remains underexplored.^76–78^ In addition, many conjugates of bile acids were also only recently discovered.^63^ Bacterial *asst* genes are known to be present in *Escherichia coli*, *Phocaeicola vulgatus* (formerly *Bacteroides vulgatus*), *Lachnospiraceae* family members, *Sutterella* species, and other bacteria.^79–83^ We found that *asst* was overall significantly more present in *ApoE*^-/-^ *FXR*^-/-^ mice (LME: *ApoE*^-/-^ IHC vs Air p = 0.240, *ApoE*^-/-^ *FXR*^-/-^ IHC vs Air p = 0.139, Air *ApoE*^-/-^ *FXR*^-/-^ vs *ApoE*^-/-^ p = 0.020, IHC *ApoE*^-/-^ *FXR*^-/-^ vs *ApoE*^-/-^ p = 0.048) (**Figure 5d**), including the final timepoint (2MWW-HB: *ApoE*^-/-^ IHC vs Air p = 0.293, *ApoE*^-/-^ *FXR*^-/-^ IHC vs Air p = 0.999, Air *ApoE*^-/-^ *FXR*^-/-^ vs *ApoE*^-/-^ p = 0.019, IHC *ApoE*^-/-^*FXR*^-/-^ vs *ApoE*^-/-^ p = 0.049) (**Figure 5e**). Based on the function of *asst*, we expected sulfation of cholic acid (CA). When looking at the last timepoint (TP6), we did observe a positive correlation between the prevalence of asst reads and the natural log ratio of sulfated cholic acid (CA) to CA (**Figure 5f**). This indicated that bacterial *asst* genes can contribute to the sulfation of bile acids. Since the formation of BA-sulfates increases during cholestatic diseases^84^, this metabolic pathway should be further explored.

### Multi-omics analysis shows bile acids are key mediators

Multi-omics analysis via joint-RPCA allows to examine co-occurrences of microbes (metagenomics) and metabolites (untargeted metabolomics). For this analysis, the initial baseline timepoint (TP1) was excluded, as mice were not on a HFHCt diet. The resulting joint-RPCA showed the biggest difference was between genotypes (PERMANOVA, pseudo-F = 89, p < 0.001) (**Figure 6a**). Secondarily, air exposure also showed significant differences within genotype (PERMANOVA, IHC vs Air: *ApoE^-/-^* pseudo-F = 28, p = 0.001; *ApoE^-/-^ FXR^-/-^* pseudo-F = 21, p = 0.001). In general, the predicted metabolic classes with the most co-occurrences with bacteria of interest present were cholane steroids (*i.e.,* bile acids) (**Figure S4**). Upon examination of the 52 bile acids that were found to be significantly altered in *ApoE^-/-^*IHC and the patterns of microbial co-occurrences (**Figure S5**), we observed a unique cluster linked to the genera *Ruthenibacterium* (**Figure 3a**), *Akkermansia* (**Figure S2a**), *Blautia A* (**Figure S2e**), and *Parasutterella* (**Figure S2f**). These connections were further explored using a network plot (**Figure 6b**). The *Blautia* genus had the most co-occurences with the bile acids of interest. This was expected as several species are known to have both *bsh* and *hsdh* genes.^85–87^ *Akkermansia*, *Ruthenibacterium*, and *Parasutterella* genera also appeared to be genera important in distinguishing *ApoE^-/-^* mice under IHC conditions from the other experimental groups. Moreover, *Ruthenibacterium* (**Figure 3a**) appeared to have a mostly positive co-occurrence with several different key bile acids, although none of them appeared to be directly correlated with *hsdh* levels. While *Ruthernibacterium intestinale* is known to encode a *bsh* gene^86^, *Ruthenibacterium lactatiformans* (**Figure 3a**) is less invetigate species, but known to be bile tolerant^88^. However, its ability to interact with or modify bile acids remains unknown. We observed 5 bile acids that have strong correlations with the *hsdh* gene (**Figure 6b**, subsets). A putative pentahydroxylated bile acid (*m/z* 498.3441, RT 6.81) and putative C27 tetrahydroxy bile alcohol sulfate (*m/z* 401.3413, RT 7.38) had positive correlations (slopes) with *hsdh* gene, whereas a putative C27 pentahydroxy bile alcohol sulfate (*m/z* 417.3359, RT 6.11), a putative trihydroxylated bile acid (*m/z* 436.3404, RT 6.19), and a putative dihydroxylated bile acid conjugated with methyl cysteamine (*m/z* 466.3321, RT 5.52) had negative correlations. Since host-produced bile acids are typically trihydroxylated, we expect microbial communities enriched in *hsdh* genes, such in the case of *ApoE^-/-^* IHC, to add or subtract hydroxyl groups and have more deviation from a steady state. We did observe both strong negative and positive correlations with such bile acids (**Figure 6b-c**). The combined effects of higher detection of *hsdh*, an increase in key bile acids (those with positive slopes outweighing those with negative slopes), and a greater burden of aortic lesions showed a moderate to strong correlation (Pearson, p<0.05 for all axes comparisons, correlation coefficient > 0.4 for all axes comparisons) (**Figure 6d**).

**Figure 6.**
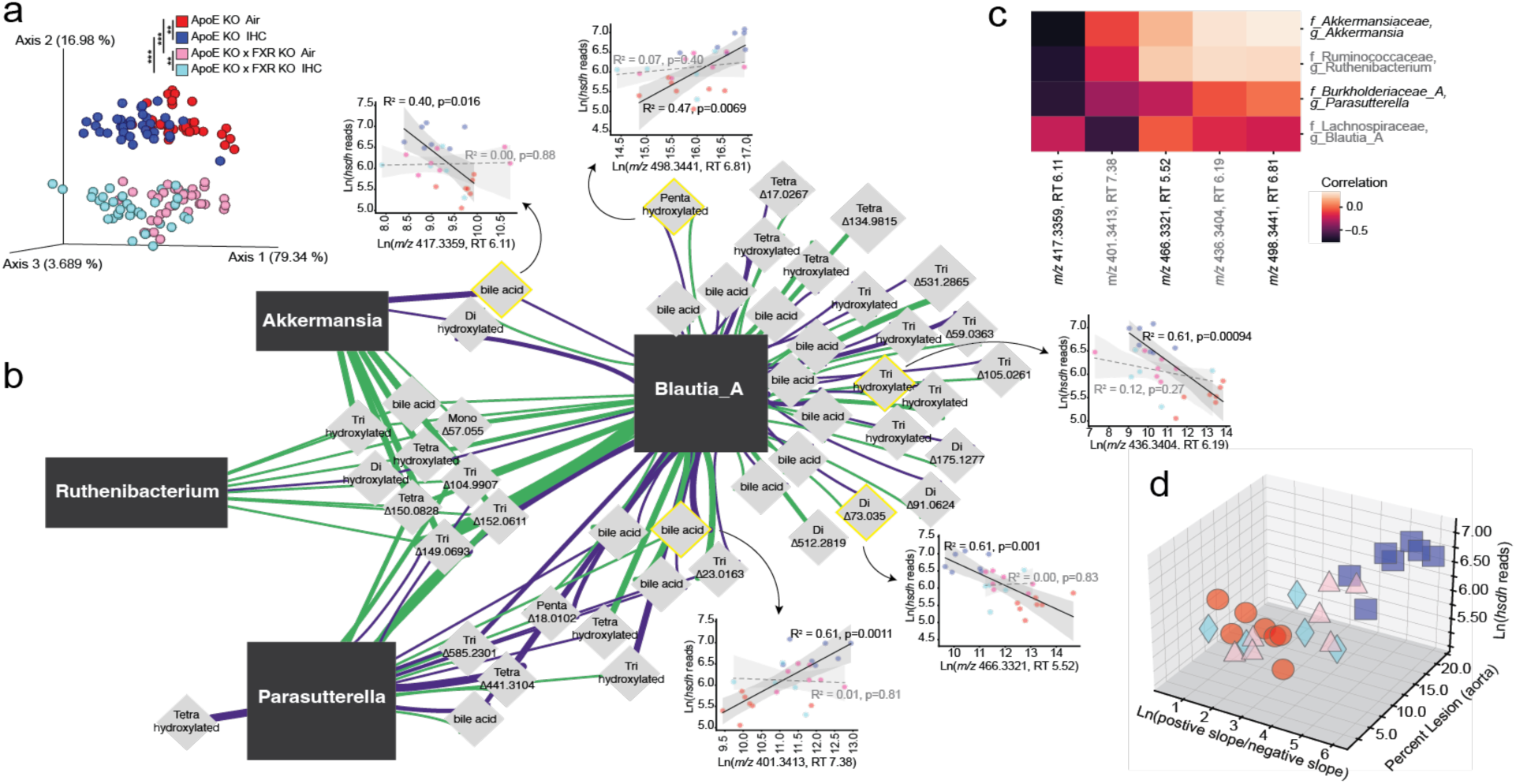
Multiomics using joint-RPCA. **a)** Joint-RPCA EMPeror plot. Significance determined by pairwise PERMANOVA with post-hoc Dunn’s test. **b)** Joint-RPCA network plot of the 52 bile acids of interest that were significantly altered in *ApoE^-/-^* IHC compared to other groups and the highlighted genera from the clustering pattern seen in **Figure S5**. Dark grey rectangles indicate bacterial genera, light grey diamonds indicate bile acids. Green indicates positive co-occurence value, purple indicates negative co-occurence value. Thickness of the lines indicates the number of repeated occurrences for OGU in that genera. Bile acids with a strong correlation to *hsdh* are highlighted in yellow and linear regression plots are shown nearby. **c)** Joint-RPCA co-occurrence clustermap of the selected genera and highlighted bile acids in **b** that strongly correlate with *hsdh* (either positive or negative). The median co-occurrence value for all detected OGU in that genus is used. And **d)** 3D scatterplot of the percent lesions in the aorta, natural log of the hsdh read count matched, and the natural log ratio of the highlighted bile acids in **b**, with those with a positive slope in the numerator and negative slope in the denominator.

## DISCUSSION

Here we show that FXR signaling serves as a key host determinant linking IHC to gut microbial metabolic reprogramming and atherosclerosis. By integrating atherosclerosis phenotype with microbiome, metabolome, and functional gene analyses in *ApoE^-/-^*mice, we show that genetic ablation of FXR significantly reduces IHC-induced aortic atherosclerosis, prevents IHC-driven restructuring of the gut microbiome, and disrupts the coupling between microbial bile acid metabolism and atherosclerosis progression. These findings establish FXR as a central mediator through which IHC stress is translated into maladaptive host–microbe interactions.

IHC significantly accelerated atherosclerosis in *ApoE^-/-^*mice, consistent with our previous studies^9,13^, whereas FXR deficiency reduced aortic lesion under IHC to levels comparable to air-exposed controls. This protective effect was vessel site-specific since pulmonary artery lesions persisted in *ApoE^-/-^ FXR^-/-^* mice, indicating that the aortic and pulmonary vascular beds respond to IHC via distinct mechanisms. While the aorta appears to rely on FXR-dependent metabolic and inflammatory signaling^20,89^, pulmonary artery atherosclerosis may be driven by alternative pathways, such as direct hypoxia-induced vascular remodeling^90^, local inflammatory responses^91^, or differences in hemodynamic stress^92^. The data highlight the complexity of vascular disease and underscore the importance of considering regional vascular biology when evaluating therapeutic targets.

Specific microbial taxa emerged as associated with atherosclerosis progression under IHC, particularly during later stages of disease. Enrichment of *Ruthenibacterium lactatiformans*, *Sutterella wadsworthensis*, *Parasutterella*, along with depletion of *Blautia A*, *Akkermansiaceae* and *Muribaculaceae*, points to coordinated changes in bile acid metabolism and gut barrier–associated taxa. For example, *Blautia* species harbor bile salt hydrolase activity^93,94^ and are closely related to human bile acid 7α-dehydroxylating taxa^95^, linking them to bile acid metabolism and metabolic homeostasis. Their abundance has been associated with improved glucose and lipid metabolism and reduced inflammation.^93,94,96,97^ *Ruthenibacterium lactatiformans*, an anaerobic, lactate-producing member of the *Ruminococcaceae* family^88^, lacks known bile acid–transforming enzymes. However, increased abundance has been reported in association with cardiovascular risk^98^, fasting^99^, and higher circulating high-density lipoprotein levels^100^, suggesting a potential role as a metabolic modifier of host lipid metabolism rather than a direct mediator of bile acid biotransformation. Both *Akkermansiaceae* and *Muribaculaceae* strengthen gut barrier integrity and constrain metabolite translocation under hypoxic stress.^13,101–105^ These changes were largely absent in *ApoE^-/-^ FXR^-/-^* mice, supporting a role for FXR in shaping a disease-permissive gut environment under IHC–i.e. gut barrier dysfunction, altered luminal bile acid composition, and increased host exposure to microbially modified metabolites–that potentially promote atherosclerosis.

Untargeted metabolomics revealed profound metabolic remodeling under IHC, with bile acids as the dominant metabolite class associated with atherosclerosis. A bile acid log-ratio signature, defined by bile acids increased versus depleted under IHC, strongly correlated with aortic lesions and was established rapidly after exposure onset and remained stable throughout the study in *ApoE^-/-^* mice but not in *ApoE^-/-^ FXR^-/-^* mice, indicating that FXR is required for bile acid remodeling to generate a permissive metabolic state for atherogenesis. Together, these data support a model in which FXR functions as a critical gatekeeper controlling the pathogenic transmission of microbially derived bile acid signals to host metabolic and inflammatory pathways.

Consistent with this notion, we identified microbial hydroxysteroid dehydrogenase (*hsdh*) genes as key mediators linking microbiome to atherosclerosis. The prevalence of *hsdh* was increased under IHC in *ApoE^-/-^*mice and correlated strongly with aortic lesion degree, but this association was lost in *ApoE^-/-^FXR^-/-^* mice. Because *hsdh* enzymes catalyze reversible oxidation–reduction of hydroxyl groups on steroid backbones, most notably bile acids^106^, increased *hsdh* activity would be expected to diversify the bile acid pool and alter FXR ligand availability. Our data implicate microbial bile acid modification by *hsdh* as a potential functional driver of FXR-dependent atherosclerosis under IHC. In contrast, *bai* and *bsh* genes, two other major bile acid modification enzymes^15,107^, were not significantly altered. This functional specificity highlights the importance of moving beyond taxonomic profiling to focus on microbial metabolic capacity. In addition, we observed increased prevalence of microbial aryl-sulfate sulfotransferase (*asst*) genes in *ApoE^-/-^ FXR^-/-^* mice, along with positive correlations between *asst* abundance and sulfated bile acid ratios. Bile acid sulfation is thought to facilitate detoxification and excretion^15,84,107^, and its enrichment in FXR deficient mice may represent a compensatory microbial response that limits bioactive bile acid signaling. These observations raise the possibility that distinct microbial bile acid modification routes, redox versus sulfation, have opposing effects on host FXR signaling and atherosclerosis susceptibility.

Integrated multi-omics analysis using joint-RPCA further demonstrated bile acids as central effectors of the host–microbe crosstalk under IHC. Co-occurrence networks revealed coordinated shifts connecting specific microbial genera, including *Blautia*, *Akkermansia*, *Ruthenibacterium*, and *Parasutterella,* with disease-associated bile acid profiles. These linkages were disrupted by FXR deficiency, emphasizing FXR’s role in shaping the interactions among the microbiome, microbially derived metabolites and atherosclerosis.

Taken together, our findings support a model in which IHC reprogram microbial bile acid metabolism and generate an altered bile acid pool that engages FXR to drive downstream metabolic and inflammatory processes leading to atherosclerosis. Thus, FXR acts as a gatekeeper that determines whether microbial bile acid signals propagate to vascular pathology. This work provides mechanistic insight into how stressors such as sleep-disordered breathing restructure microbial function to influence host metabolism and systemic disease. Targeting FXR signaling or specific microbial bile acid–modifying pathways may offer therapeutic strategies to mitigate hypoxia-associated cardiometabolic disease.

## ACKNOWLEDGEMENTS

This study was supported by the National Institutes of Health grant R01 HL157445 to GGH. This publication includes data generated at the UC San Diego IGM Genomics Center utilizing an Illumina X Plus that was purchased with funding from a National Institutes of Health SIG grant (#S10 OD026929).

## AUTHOR CONTRIBUTIONS

R.K. and G.G.H. conceived and designed research; O.P. and J.M. performed mouse experiments; J.X., O.P., and D.Z. analyzed physiology data; C.A. and D.M. analyzed microbiome and multi-omics data; S.Z. analyzed metabolome and multi-omics data. J.X., C.A., S.Z., P.D., R.K., and G.G.H. interpreted results of experiments; J.X., C.A., S.Z., D.Z., and O.P. prepared figures; J.X., C.A., S.Z. and D.Z. drafted manuscript; All authors edited, revised, and approved the manuscript.

