## Supplementary Figure 1 for "Farnesoid X receptor-dependent microbiome-bile acid signaling mediates obstructive sleep apnea-induced atherosclerosis"

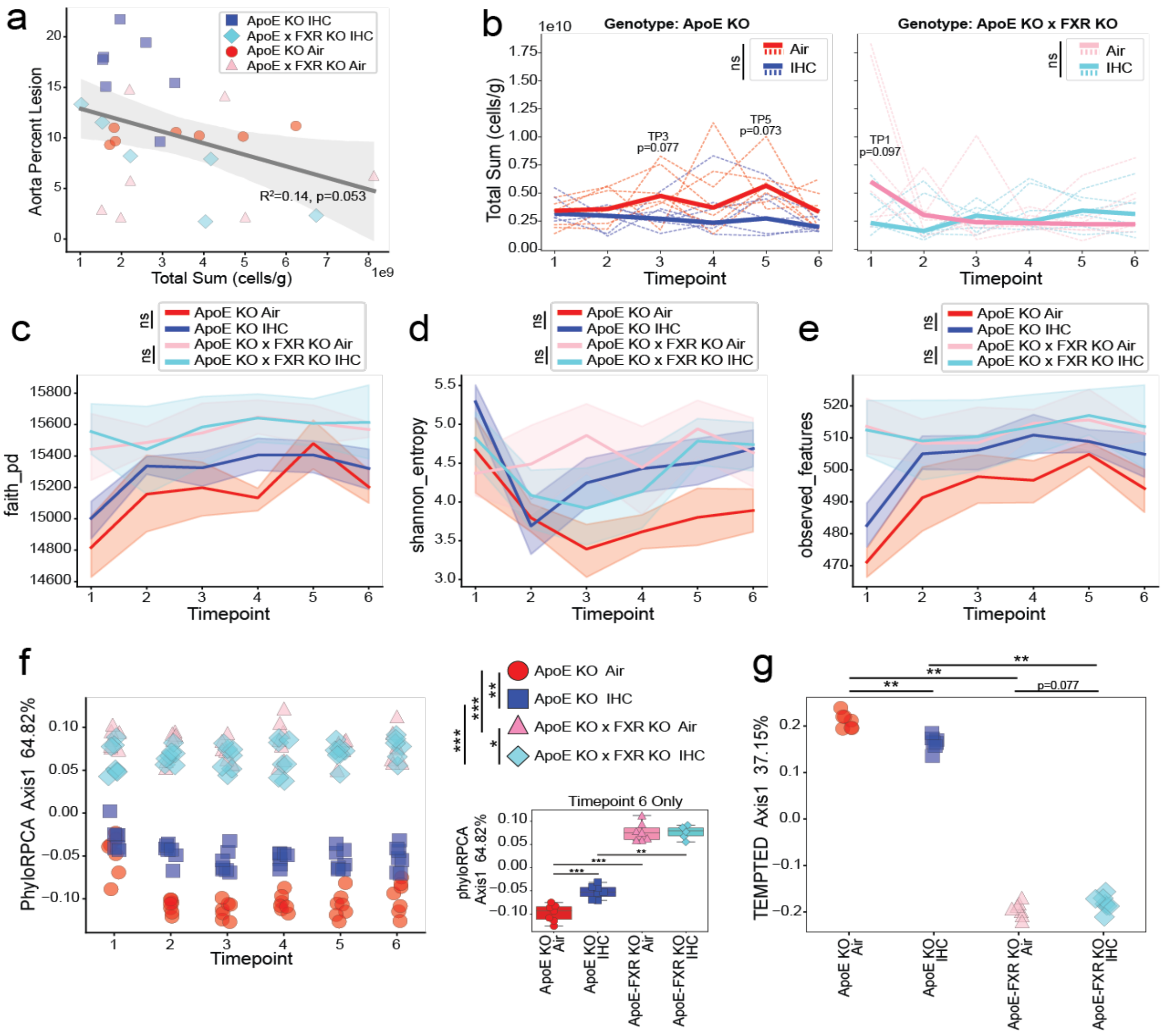


**Supplemental Figure 1.** **Additional Alpha and Beta Diversity Plots. a)** Absolute abundance (cells/g) comparison to percent lesions in aorta at the final timepoint only (TP6). **b)** Absolute abundance (cells/g) comparison by genotype and exposure across all timepoints. Individual lines represent a unique host, thicker dashed lines indicate median value. Significance determined by linear mixed effects model (LME). Then, lineplots of alpha diversity metrics **c)** Faith's PD, **d)** Shannon, and **e)** observed OGU by genotype and exposure; significance determined by LME. Shaded regions indicate 95% confidence intervals. **f)** Beta diversity metric phyloRPCA Axis 1 across the timepoints; significance determined by pairwise PERMANOVA with post-hoc Dunn's test. Subset boxplot/stripplot is for TP6 only; significance determined by two-sided Mann-Whitney-Wilcoxon with Benjamini-Hochberg correction. And **g)** longitudinal beta diversity metric TEMPTED Axis 1 by genotype and exposure. Significance determined by pairwise PERMANOVA with post-hoc Dunn's test. Notation: * $p < 0.05$, ** $p < 0.01$, *** $p < 0.001$, **** $p < 0.001$.
