## Supplementary Figure 2 for "Farnesoid X receptor-dependent microbiome-bile acid signaling mediates obstructive sleep apnea-induced atherosclerosis"

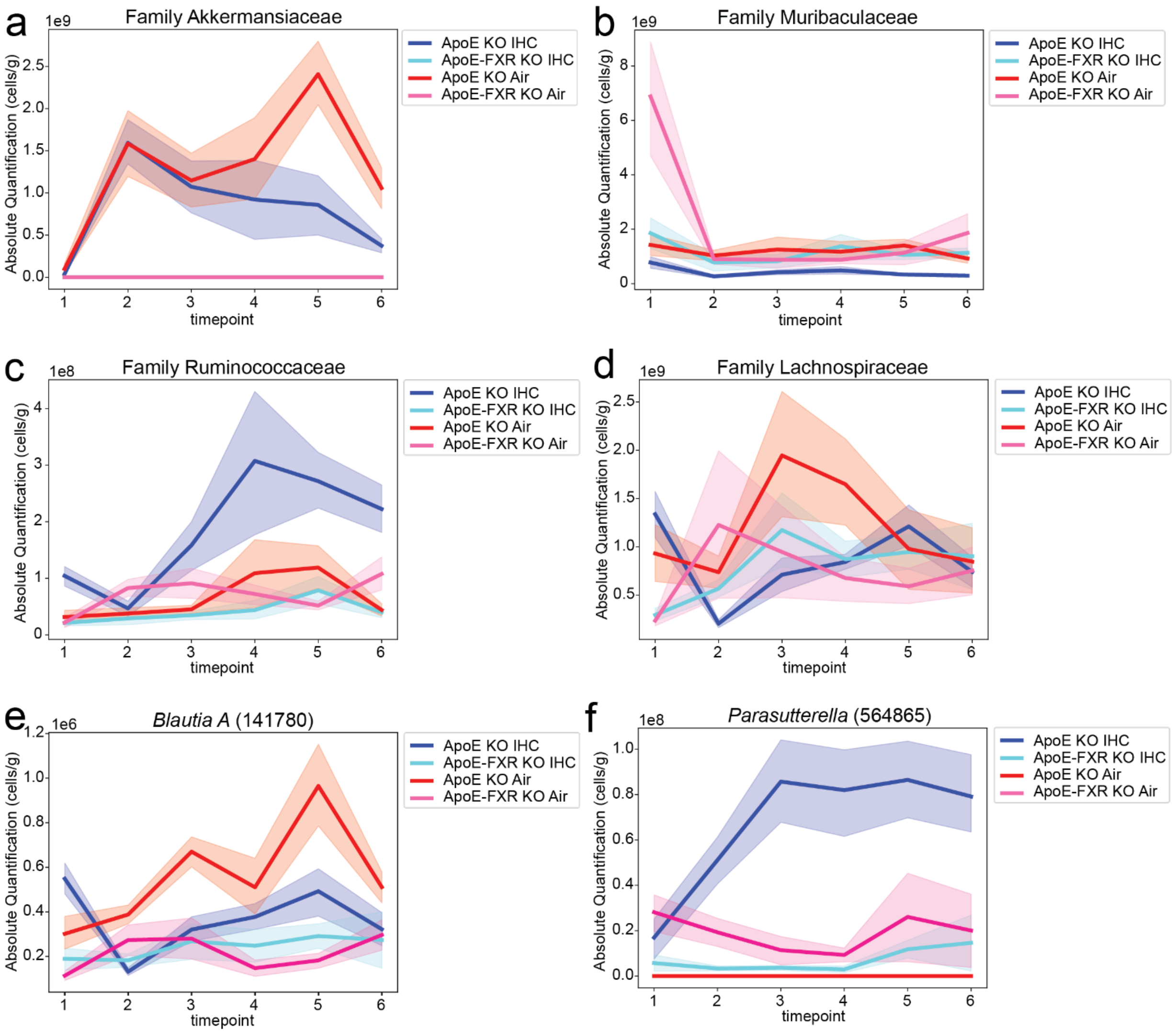


**Supplemental Figure 2. Additional Relevant Family Level absolute abundance (cells/g) Plots.** Lineplot of *Akkermansiaceae* (**a**), *Muribaculaceae* (**b**), *Ruminococcaceae* (**c**), *Lachnospiraceae* (**d**), *Blautia A* (141780) genus [*Lachnospiraceae*] (**e**), and *Parasutterella* genus [*Burkholderiaceae* A] (**f**) by genotype and exposure over time. Solid line indicates the mean and the shaded region indicates standard error of the mean (SEM).
