## Supplementary Figure 3 for "Farnesoid X receptor-dependent microbiome-bile acid signaling mediates obstructive sleep apnea-induced atherosclerosis"

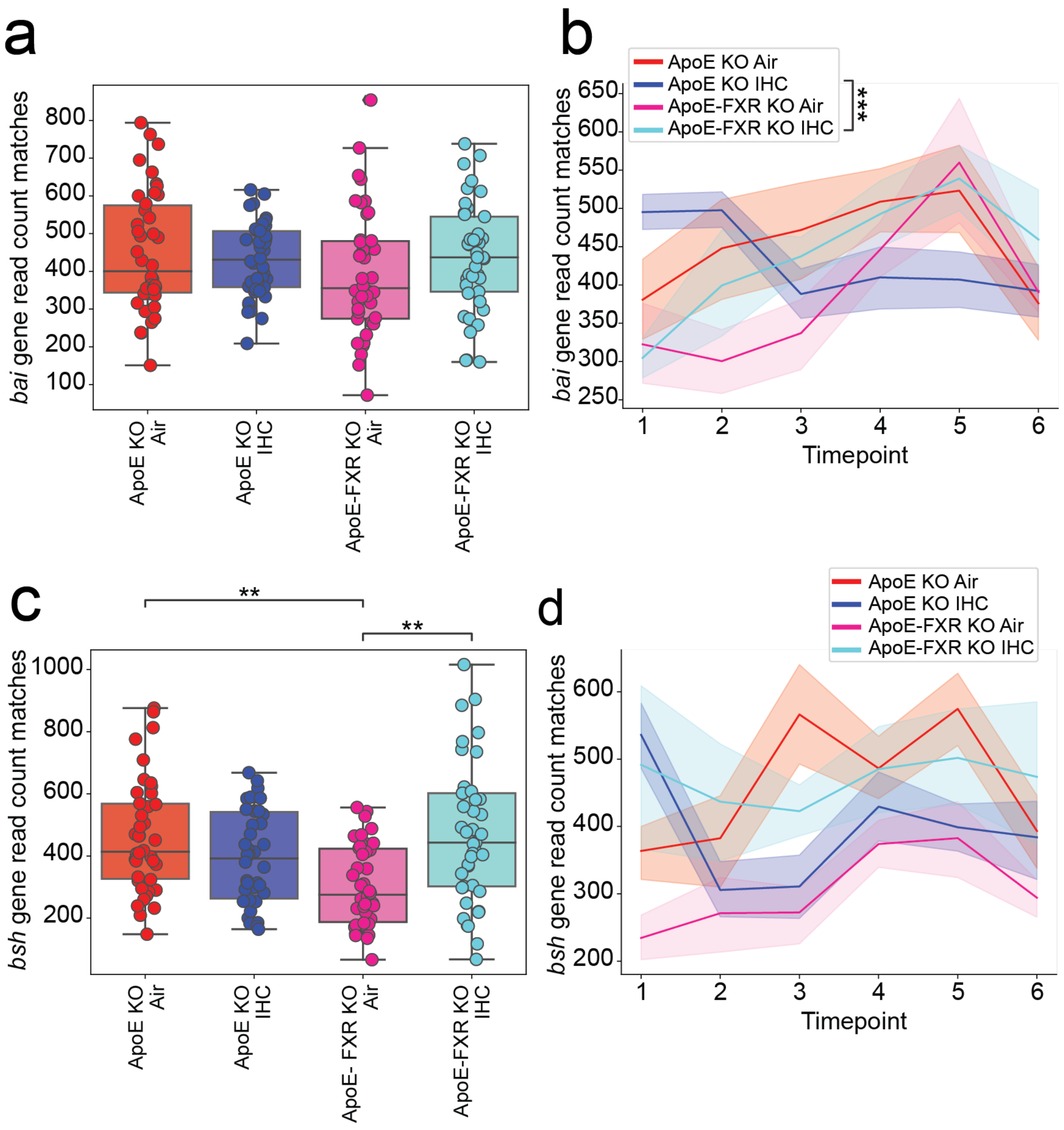


**Supplemental Figure 3.** **Additional Bile Acid Modification Enzymes. a)** Box-scatterplot of *bai* read count matches by genotype and exposure. No significant comparisons found. **b)** Lineplot of *bai* read count matches by genotype and exposure over time; shaded region indicates standard error of the mean (SEM). **c)** Box-scatterplot of *bsh* read count matches by genotype and exposure. And **d)** lineplot of *bsh* read count matches by genotype and exposure over time. No significant comparisons found. For lineplots, the shaded region indicates standard error of the mean (SEM). Significance for a, c-f is determined by LME and accounts for repeated measures; lineplots show significance for interaction term. Notation: * $p < 0.05$, ** $p < 0.01$, *** $p < 0.001$.
