## Supplementary Figure 4 for "Farnesoid X receptor-dependent microbiome-bile acid signaling mediates obstructive sleep apnea-induced atherosclerosis"

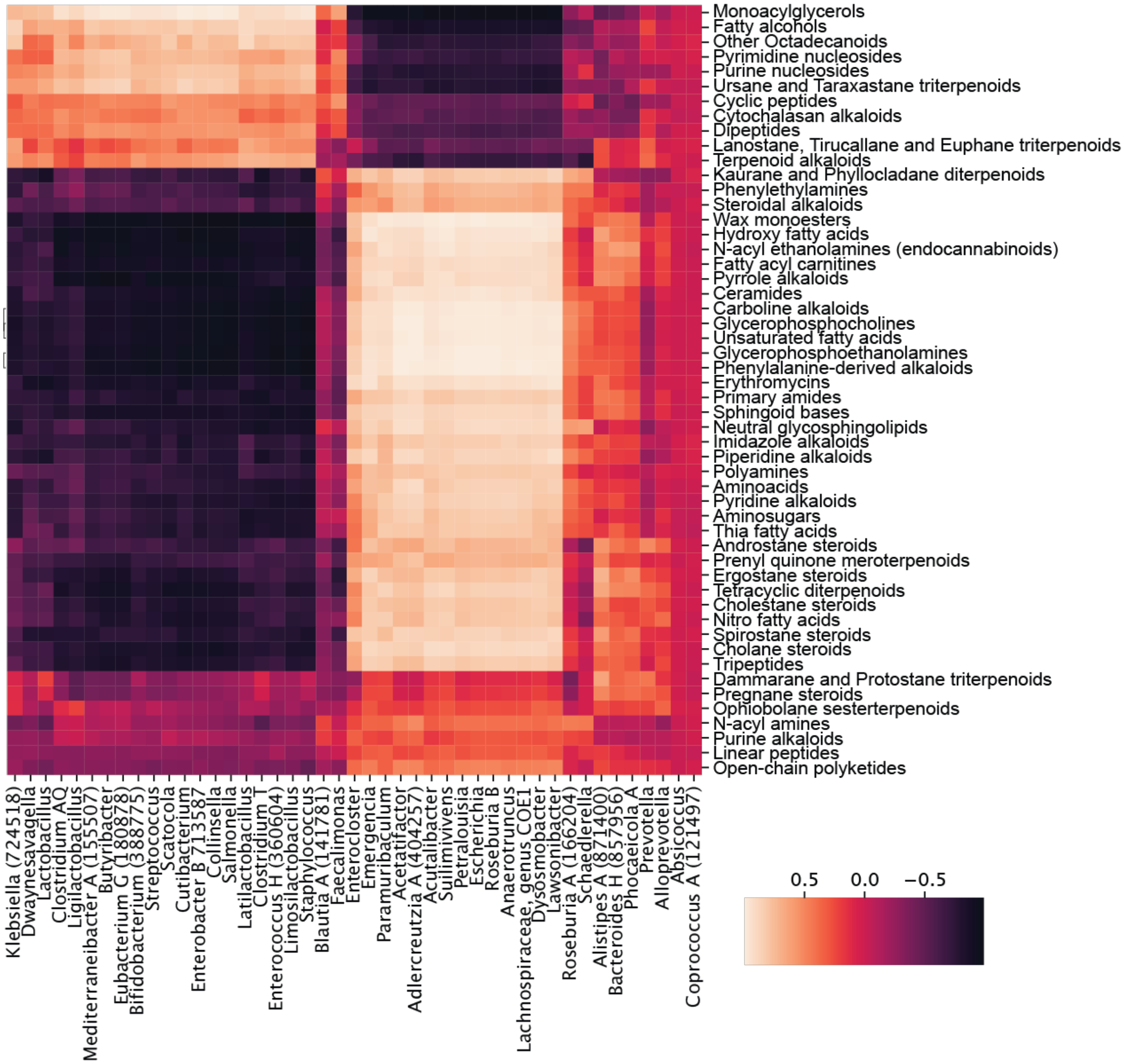


**<u>Supplemental Figure 4.</u> Multiomics, top genera and metabolite classes.** Joint-RPCA co-occurrence cluster map with the top most prevalent genera and predict metabolite classes via CANOPUS. The top genera are those that had more than 5000 co-occurrences with values greater than 0.75 absolute value (both positive and negative). The top metabolite classes are those that had more than 1000 co-occurrence values greater than 0.75 absolute value. Grouped by genus and NPC Class, median co-occurrence value is shown.
