## Supplementary Figure 5 for "Farnesoid X receptor-dependent microbiome-bile acid signaling mediates obstructive sleep apnea-induced atherosclerosis"

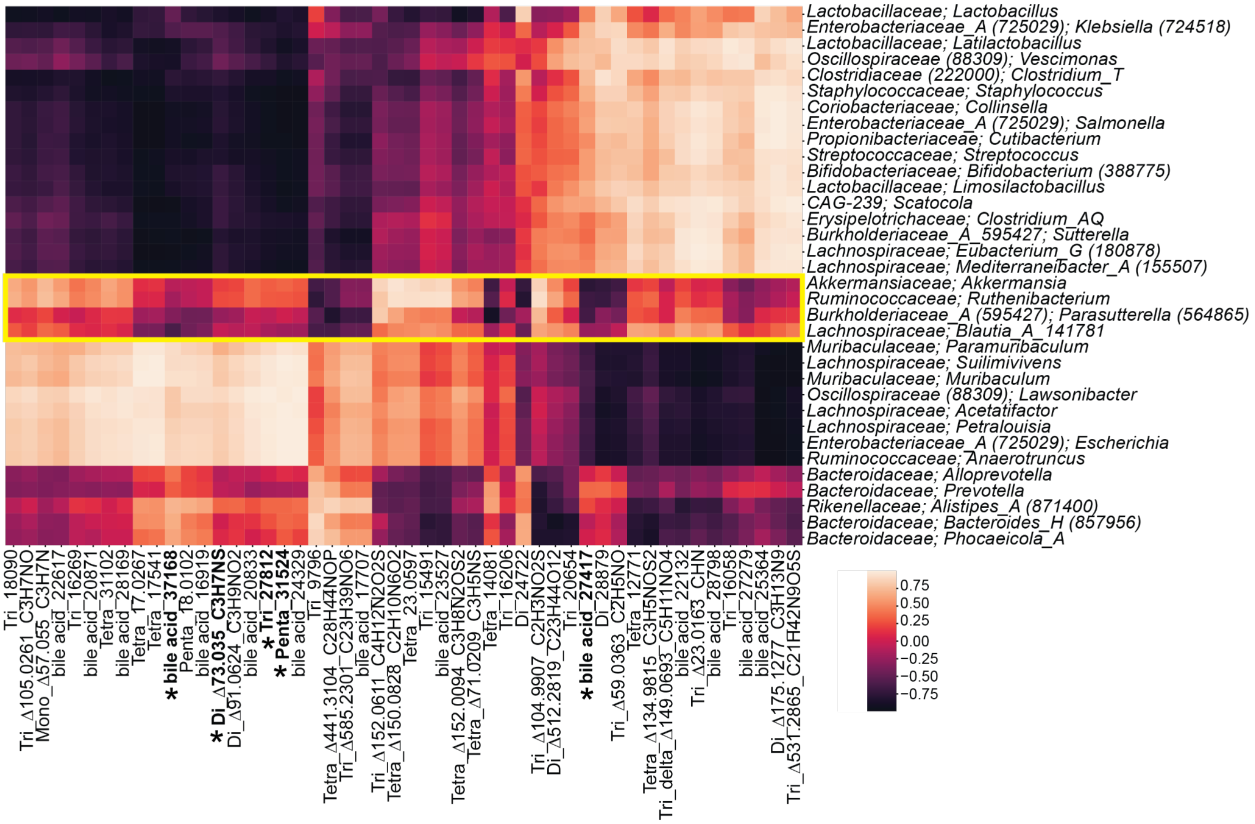


**<u>Supplemental Figure 5.</u> Multiomics, top genera that co-occur with bile acids of interest.** Joint-RPCA co-occurrence clustermap with the top genera connected to the 52 bile acids of interest classified by number of hydroxylated carbons (or simply listed as bile acid if putative classification without diagnostic ions) - that had more than 5 co-occurrences with values greater than an absolute value of 0.75 plus *Sutterella*, *Parasutterella*, *Ruthenibacterium*, *Akkermansia*, and *Muribaculum*, which were found to be important in other analyses (Figure 3, S2). The 52 bile acids of interest were determined by modelling the effects of IHC and association with atherosclerosis across [ApoE$^{-/-}$] and [ApoE$^{-/-}$; FXR$^{-/-}$]. The yellow box indicates a cluster of interest. Bile acids that had a correlation with *hsdh* in ApoE$^{-/-}$ were bolded and marked by star, also visualized in Figure 6b.
